# Inactivated voltage-gated sodium channels are preferentially targeted in cinacalcet block of mouse neocortical action potentials

**DOI:** 10.1101/2022.08.15.504002

**Authors:** Jamie S. Lindner, Salil R. Rajayer, Stephen M. Smith

## Abstract

Voltage-gated sodium channel (VGSC) activation is essential for action potential generation in the brain. Allosteric calcium-sensing receptor (CaSR) agonist, cinacalcet, strongly and ubiquitously inhibits VGSC currents in neocortical neurons via an unidentified, G-protein-dependent blocking molecule. The mechanisms by which VGSC characteristics influence cinacalcet-mediated inhibition are not well understood. Here, using whole-cell patch clamp methods, we investigated the voltage-dependence of cinacalcet-mediated inhibition of VGSCs and the channel state preference of cinacalcet. The rate of inhibition of VGSC currents was accelerated at more depolarized holding potentials. Cinacalcet shifted the voltage-dependence of both fast and slow inactivation of VGSCs in the hyperpolarizing direction. Utilizing a simple model, the voltage-dependence of VGSC current inhibition may be explained if the affinity of the blocking molecule to the channel states follows the sequence: fast-inactivated > slow-inactivated > resting. The state dependence of block contributes to the non-linearity of action potential block by cinacalcet. This dynamic and abundant signaling pathway by which G-proteins regulate VGSC currents provides an important voltage-dependent mechanism for modulating central neuronal excitability.

**Key points summary:** Voltage-gated sodium channels are essential for the action potential generation and propagation that is central to physiological function in excitable cells. VGSC inhibitors are useful therapies to treat epilepsy, pain, and cardiac arrhythmias.

Cinacalcet inhibits VGSC currents strongly and the underlying G-protein dependent signaling pathway occurs in the vast majority of neocortical and hippocampal neurons.

Here we demonstrate that cinacalcet inhibits the VGSC current by activating a downstream blocking molecule that preferentially binds to the fast-inactivated state, that the blocking molecule stabilizes the inactivated states, and that cinacalcet impacts neuronal excitability in a non-linear manner.

Characterization of the mechanism by which cinacalcet operates will facilitate the determination of its role in regulating neocortical excitability and the identification of new therapeutic targets for epilepsy, pain, and arrythmias.

## Introduction

Voltage-gated sodium channels (VGSC) are essential for the action potential generation and propagation that is central to physiological function in excitable cells (Hodgkin & Huxley, 1952; Hille, 2001). VGSCs are targets for a wide range of important drugs used as local anesthetics, antiarrhythmics, and anticonvulsants and operate by reducing excitability in cardiac and central nervous tissue (Catterall, 1999; Hille, 2001; Nau & Wang, 2004).

Specificity of action of different VGSC inhibitors across cell types arises from a number of factors. Tissue-specific variation in VGSC subtypes is an important contributor to the selectivity of effect since VGSC modulator affinity may vary with VGSC subunit isoform (Mantegazza *et al*., 2005; England & de Groot, 2009; Okura *et al*., 2014). Preferential binding to specific channel states may also influence the tissue specific effectiveness of the inhibitor. If the inhibitor binds preferentially to a particular channel state, it will be more effective in tissues with membrane potentials that increase the fraction of channels in that state (Kuo & Bean, 1994; Karoly *et al*., 2010; Theile *et al*., 2016). If the VGSC inhibitor operates indirectly via a G-protein coupled receptor (GPCR), this also confers selectivity of effect (Cantrell *et al*., 1996; Mattheisen *et al*., 2018). Regional specificity may arise from variable expression of the GPCR or the downstream signaling pathway, as both will determine if cells are modulated by the VGSC inhibitor. The degree of VGSC inhibition by G-protein regulation ranges from 10-100% depending on brain regions and GPCR identity (Cantrell *et al*., 1996; Cantrell *et al*., 1997; Carr *et al*., 2002; Carlier *et al*., 2006; Mattheisen *et al*., 2018). Cinacalcet, a calcium-sensing receptor (CaSR) allosteric agonist, strongly and almost ubiquitously inhibited VGSC currents via an unidentified, indirect pathway mediated by G-proteins but independent of the CaSR (Mattheisen *et al*., 2018). The high abundance and strength of inhibition of this pathway positions it to have an important impact on neuronal excitability.

To better understand how neuronal excitability is affected by the pathway utilized by cinacalcet to inhibit VGSC currents, we studied how VGSC properties are affected following the application of this drug to neocortical neurons. Here, we show that cinacalcet activity is enhanced at more depolarized holding potentials, indicating a preference of an unidentified downstream blocking molecule (X) for the inactivated state. Reversal of cinacalcet-mediated inhibition of VGSC currents via prolonged hyperpolarization suggested that the mechanism may involve stabilization of the slow-inactivated state. We investigated the correlation between the development of inactivation and the kinetics of block by cinacalcet. The data support a model indicating that X binds to the various channel states with the preference fast-inactivated> slow-inactivated> resting state.

## Materials and Methods

### Ethical approval

All animal procedures were approved by VA Portland Health Care System Insitutitonal Animal Care and Use Committee (IRBNetID: 1635414-4) in accordance with the US Public Health Service policy on Humane Care and Use of Laboratory Animals and the National Institutes of Health Guide for the Care and Use of Laboratory Animals. Neocortical neurons were isolated from 1-2 day old postnatal mouse pups of either sex as described previously (Martiszus *et al*., 2021; Ritzau-Jost *et al*., 2021). Animals were decapitated following general anesthesia with isoflurane and cerebral cortices were removed. Cortices were incubated in trypsin and DNase and dissociated with heat polished pipettes. Dissociated cells were cultured in MEM plus 5% FBS on glass coverslips. Cytosine arabinoside (4 μM) was added 48-72 hours after plating to limit glial division. Cells were used, unless otherwise stated, after 7-35 days in culture.

### Electrophysiological Recordings

Cells were visualized with an inverted microscope (Leica DM IRB or Olympus IX70). Whole-cell voltage-clamp recordings were made from cultured neocortical neurons using an Axopatch 200B amplifier. Current clamp recordings were made using a Heka EPC10 amplifier. The preparation was continously perfused with solution which contained (in mM) 150 NaCl, 4 KCl, 10 HEPES, 10 glucose, 1.1 MgCl_2_, 1.1 CaCl_2_, pH 7.35 with NaOH. Synaptic transmission was blocked by the addition of (in μM) 10 CNQX, 10 Gabazine, and 50 APV in extracellular bath solution. In current clamp recordings, 2 mM CsCl was added to the extracellular bath solution to reduce contributions of HCN. Voltage-clamp recordings were made using a caesium methanesulfonate intraceullar solution containing (mM) 135 caesium methansulfonate, 1.8 EGTA, 10 HEPES, 4 MgCl_2_, 0.2 CaCl_2_, 0.3 NaGTP, 4 NaATP, 14 phosphocreatine disodium, pH 7.2 with TEA hydroxide. Current-clamp recordings were made using potassium-gluconate containing intracellular solution containing (mM) 135 Potassium gluconate, 10 HEPES, 4 MgCl_2_, 0.3 NaGTP, 4 NaATP, 10 phosphocreatine disodium, pH 7.2 with potassium hydroxide. Electrodes used for recording had resistance ranging from 3 to 8 MΩ. Voltages have been corrected for liquid junction potentials. All experiments were performed at room temperature (21-23 °C).

### Data Acquisition and Analysis

Whole cell voltage-clamp recordings were made using an Axopatch 200B Amplifier, filtered at 5 kHz using a Bessel filter, and sampled at 20 kHz during acquisition. Whole cell current-clamp recordings were made using a Heka EPC 10 amplifier, filtered at 2.9 kHz using a Bessel filter and sampled at 20 kHz during acqusition. Series resistance compensation was performed manually prior to acquisition. Analysis was performed using Igor Pro (Wavemetrics, Lake Oswego, OR). Data values were reported as mean (± SEM) or median, if not normally distributed. Statistical significance was determined with appropriate parametric or non-parametric tests (Excel, Graphpad prism or Igor Pro).

### Solution Application

Solutions were applied by gravity from a glass capillary (1.2 mm outer diameter) placed 1-2 mm from the neuron under study. Solutions were switched manually using a low dead volume manifold upstream of the glass capillary. CNQX and Gabazine were supplied by Abcam. Creatine Phosphate was supplied by Santa Cruz Biotech. Cinacalcet was supplied by Toronto Research Chemicals. All other reagents were supplied by Sigma-Aldrich.

## Results

### Voltage-dependence of cinacalcet-induced inhibition of VGSCs

Most pharmacological modulators of VGSC currents act directly on the ion channel and inhibit the sodium currents responsible for action potential generation. Cinacalcet is unusual because it acts indirectly, via a pathway mediated by G-proteins, to block VGSC currents (Mattheisen *et al*., 2018). We tested if the pathway preferentially interacts with specific VGSC states by comparing the effect of holding potential, on the rate of cinacalcet-induced block of VGSCs. In voltage clamp recordings, VGSC currents were elicited by test pulses to −20 mV (at 0.5 Hz) in cultured neocortical neurons perfused with Tyrode solution (containing 10 μM CNQX, 50 μM APV, and 10 μM Gabazine to block glutamatergic and GABAergic activity). After establishing a stable VGSC current baseline, 5 μM cinacalcet was applied to the neuron which reduced VGSC current amplitude (Figure 1). The time course of inhibition of VGSC currents by cinacalcet was described by a squared exponential function:

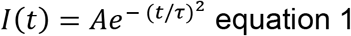

where I (t), A, t, and *τ* represent VGSC current amplitude after application, initial VGSC current amplitude, time, and time constant respectively (Mattheisen *et al*., 2018). In contrast, directly-acting inhibitors usually produce a single exponential pattern of block (Figure 1C;(Bean *et al*., 1983; Jo & Bean, 2017). The rate of block was faster at depolarized holding potentials as illustrated by the average diary plots and confirmed both by the voltage-dependence of fractional block at 200 s and the time constant of inhibition (Figure 1B-E). Hyperpolarization of the holding potential over the range of −60 to −100 mV increased *τ* five-fold (Kruskal-Wallis test, P = 0.001; Figure 1E). VGSC block by cinacalcet was incomplete at −100 mV. The accelerated block of sodium currents by cinacalcet at depolarized holding potentials probably reflects an increased affinity of the unidentified downstream blocking molecule to inactivated states of the VGSCs. These data support the proposal that block of VGSC current by cinacalcet is voltage-dependent and occurs via an indirect pathway.

**Fig. 1.**
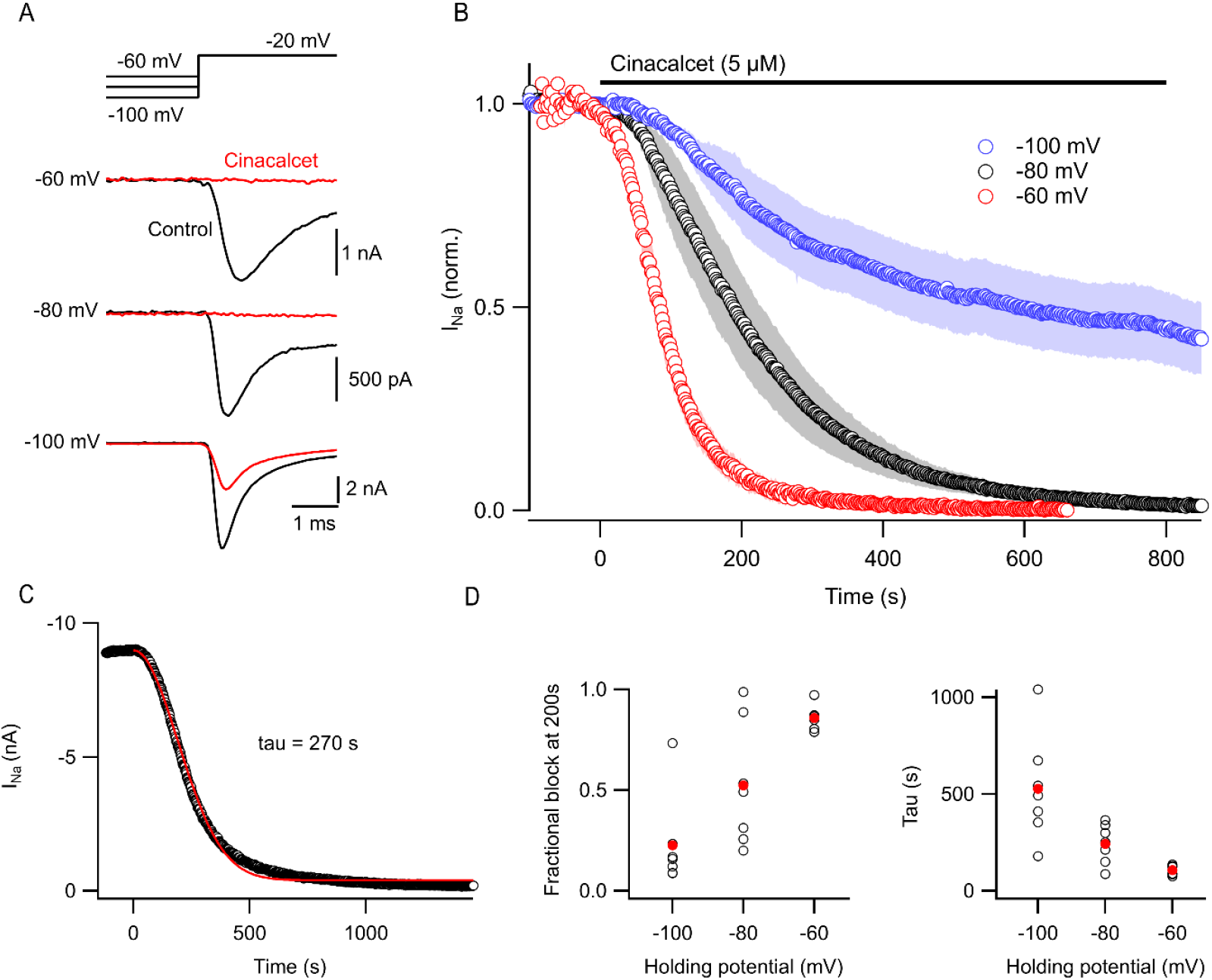
Inhibition of VGSC currents by cinacalcet is voltage-dependent. (A) Representative voltage traces show VGSC current baseline prior to (control, black) and following maximal block by 5 μM cinacalcet (red). Cells were held at −60, −80, or −100 mV and a test pulse to −20 mV was elicited every 2 s. (B) Diary plot of average normalized peak VGSC current elicited by 20 ms test pulse every 2 s from a holding potential of −60 (red, n = 6), −80 (black, n = 6), or −100 mV (blue, n = 6). Control current baseline was established prior to cinacalcet addition at time 0. (C) Exemplar diary plot of peak VGSC current elicited as in (B) following application of 5 μM cinacalcet. Data fit with squared exponential *f*/(*t*) = *A* + *Be*^−(*t*/*τ*)2^ shown in red. (D) Plot of individual (black, open circles) and median (red, filled circles) values of fractional block at 200s after application of 5 μM cinacalcet. (E) Plot of individual (black, open circles) and mean (red, filled circles) values of tau (left).

### Cinacalcet shifts the voltage-dependence of fast inactivation

Preferential binding to a specific channel state will shift the dynamic equilibrium altering the relative fraction of VGSCs that occupy each state, as explained by the modulated receptor hypothesis (Hille, 1977, 1978; Bean *et al*., 1983). Initially we tested if VGSC recovery from fast inactivation was affected by cinacalcet. The currents were elicited by a test pulse to −20 mV following a family of 100 ms prepulses (−140 to 20 mV in 10 mV increments) and these control currents were compared with those activated following ~50% or full inhibition by cinacalcet (5 μM; Figure 2A). Cinacalcet shifted the voltage-dependence of fast inactivation in the hyperpolarizing direction, with a mid-point (V_0.5_) of −58.3 ± 0.8 mV in control (n = 8), −69.4 ± 2.5 mV after half block (n = 8), and −91.9 ± 3.6 mV following full inhibition (n = 8, P = 0.00008; Figure 2B, C). The hyperpolarizing prepulses facilitated recovery of only ~10% of the fully blocked VGSC currents from inhibition by cinacalcet (Figure 2A, B). Cinacalcet also decreased the slope of the inactivation curves from −4.2 ± 0.2 mV^−1^ (control; n = 8), to −2.9 ± 0.4 mV^−1^ following half block (n = 8), and −1.9 ± 0.1 mV^−1^ following full inhibition (n = 8, P = 0.0005; Figure 2D). These data indicate that cinacalcet stabilized the fast-inactivated state of VGSCs.

**Fig. 2.**
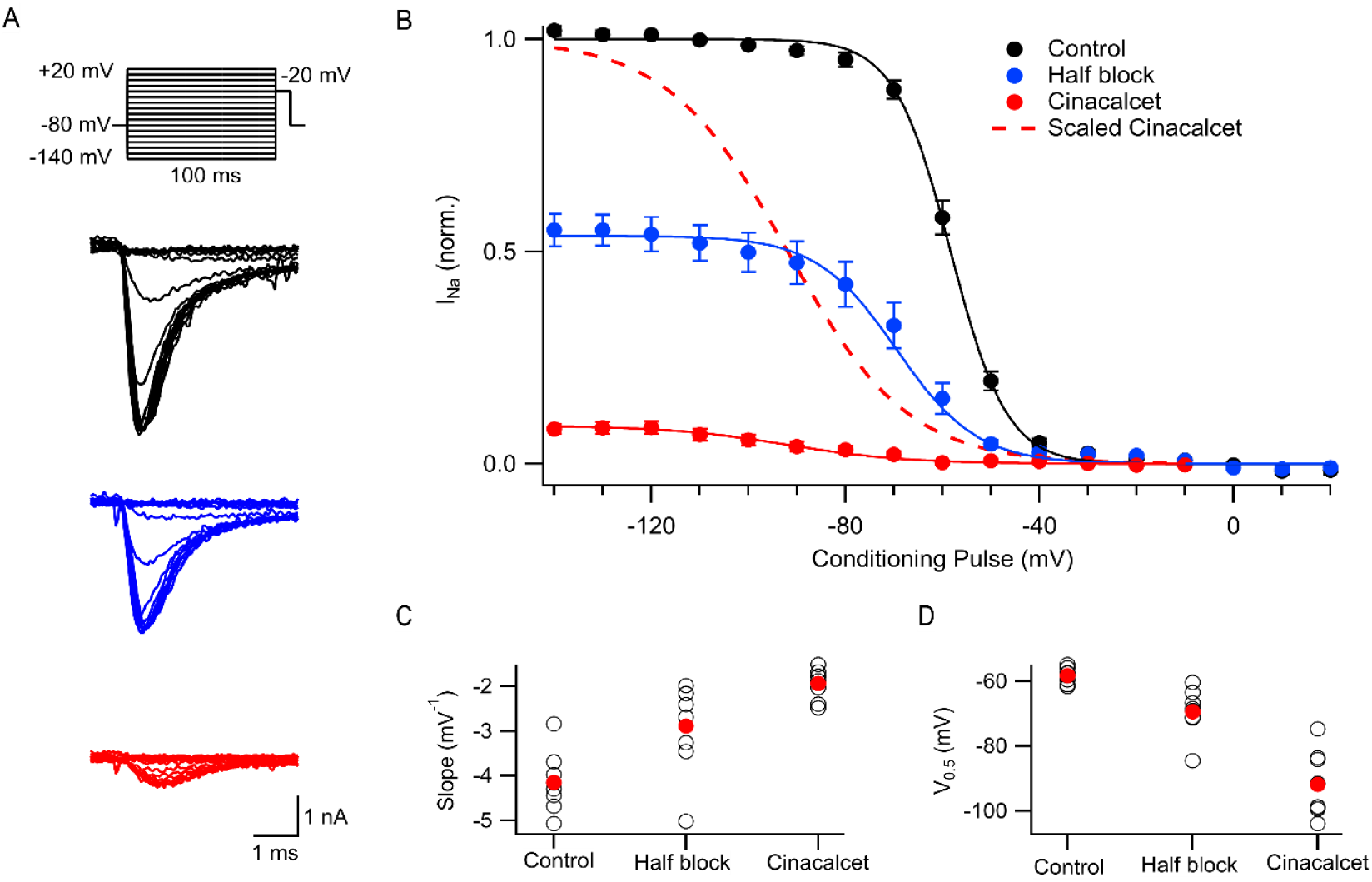
Cinacalcet shifted the voltage-dependence of fast inactivation. (A) Representative traces from a protocol used to isolate and evaluate the voltage-dependence of fast inactivation. Measurements were taken by applying a 100 ms depolarization to varying voltages (−140 to +20 mV in 10 mV increments) from a holding potential of −80 mV, and delivering a test pulse to −20 mV in control conditions (black), following inhibition of half of the starting current by 5 μM cinacalcet (blue), and following full inhibition (red). (B) Plot of average normalized peak VGSC currents elicited by test pulse following 100 ms conditioning pulse in control (black; n = 8), following inhibition of half of the starting current by cinacalcet (blue; n = 8), and following full inhibition (red; n = 8) fit to the Boltzmann equation. Dashed red line indicates average cinacalcet fit normalized to the average control maximum. (C) Plot of individual (black, open circles) and mean (red, filled circles) slope values. (D) Plot of individual (black, open circles) and mean (red, filled circles) V_0.5_ values.

### Cinacalcet stabilizes slowly recovering channel states

To further investigate the blocking mechanism of cinacalcet, we explored the possibility that X interacts with the slow-inactivated state of the VGSC. A test pulse to −20 mV, delivered after a 100 ms interval at −120 mV, evoked VGSC currents following a series of 5 s prepulses (−140 to +20 mV in 10 mV increments; Figure 3A). The 100 ms interval at −120 mV facilitated recovery from fast inactivation, so that the test pulse permitted the comparison of the effects of the prepulse on VGSC current recovery from a slow-inactivated state. Slow inactivation was complete at −10 mV in control with ~45% of VGSC currents still available for activation (Fig. 3). In the presence of 5 μM cinacalcet, we observed a substantial increase in the fraction of channels recovering slowly (96 ± 0.6%; n=6) compared to control (56 ± 4%; n=6; Figure 3B). We also observed a significant increase in the rate of slow inactivation development, with the average slopes increasing from 2.3 ± 0.2 mV^−1^ in control (n = 6) to 4.5 ± 0.5 mV^−1^ following inhibition by cinacalcet (n = 6, P = 0.004; Figure 3C). In addition, the voltage-dependence of slow inactivation shifted in the hyperpolarizing direction in the presence of cinacalcet, with a midpoint of −59.2 ± 1.7 mV in control and −97.3 ± 1.2 mV (P = 0.004) following inhibition by cinacalcet (Figure 3D). Taken together, these results indicate that cinacalcet apparently stabilizes the slow-inactivated VGSC state.

**Fig. 3.**
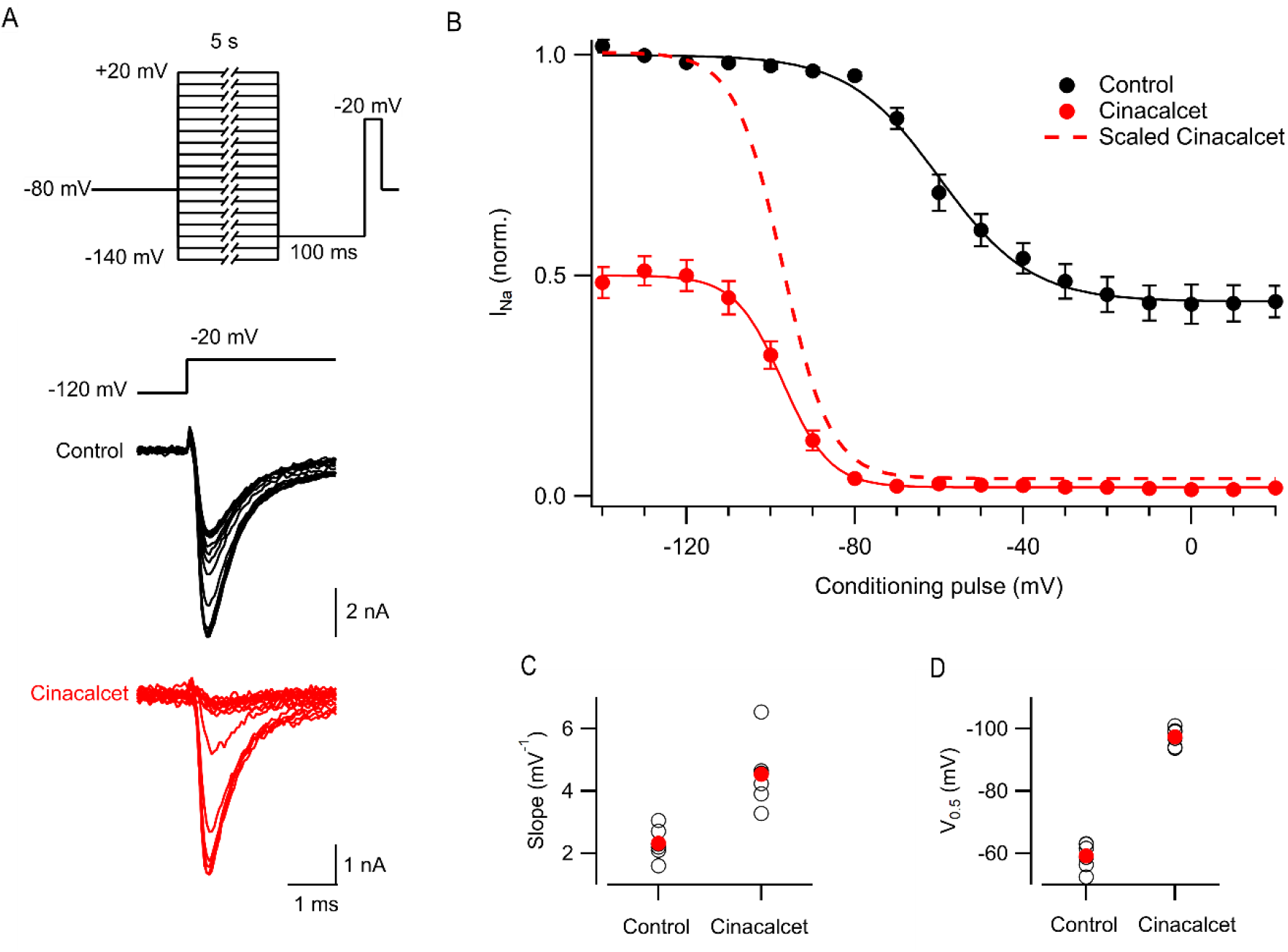
Cinacalcet enhancement of slowly recovering VGSCs. (A) Representative traces from a protocol used to isolate and evaluate the voltage-dependence of slow inactivation. Measurements were taken by applying a 5 s depolarization to varying voltages (−140 to +20 mV in 10 mV increments) from a holding potential of −80 mV, returning to −120 mV for 100 ms to allow recovery from fast inactivation, and delivering a test pulse to −20 mV in control conditions (black) and following full inhibition by 5 μM cinacalcet (red). (B) Plot of average normalized peak VGSC currents elicited by test pulse following 5 s conditioning pulse and 100 ms recovery period in control (black; n = 6) and following full inhibition by cinacalcet (red; n = 6) fit to the Boltzmann equation. Dashed red line indicates average cinacalcet fit normalized to the average control maximum. (C) Plot of individual (black, open circles) and mean (red, filled circles) slope values. (D) Plot of individual (black, open circles) and mean (red, filled circles) V0.5 values

### Cinacalcet-mediated block occurs more rapidly than slow inactivation of VGSC currents

We attempted to further characterize the kinetics of block by cinacalcet in our neocortical neurons using voltage protocols designed to differentiate between the relative affinities for fast and slow-inactivated VGSC states (Kuo & Bean, 1994; Errington *et al*., 2008). The presence of channels in the slow-inactivated state was measured with a variable length (0-16 s) prepulse to either −70, −50, −20, or +10 mV, followed by a brief recovery period at −120 mV to allow recovery from fast inactivation, and a test pulse to −20 mV (Figure 4A). Figure 4B shows the induction of slow inactivation at different voltages in the absence of cinacalcet. There was minimal slow inactivation at −70 mV; almost all VGSCs recovered from inactivation during the 50 ms recovery period even after 16 s at −70 mV. The occupancy of the slow-inactivated state increased as the inactivating pulse was made more positive, with saturation at ~93% with a 16 s pulse at +10 mV.

**Fig. 4.**
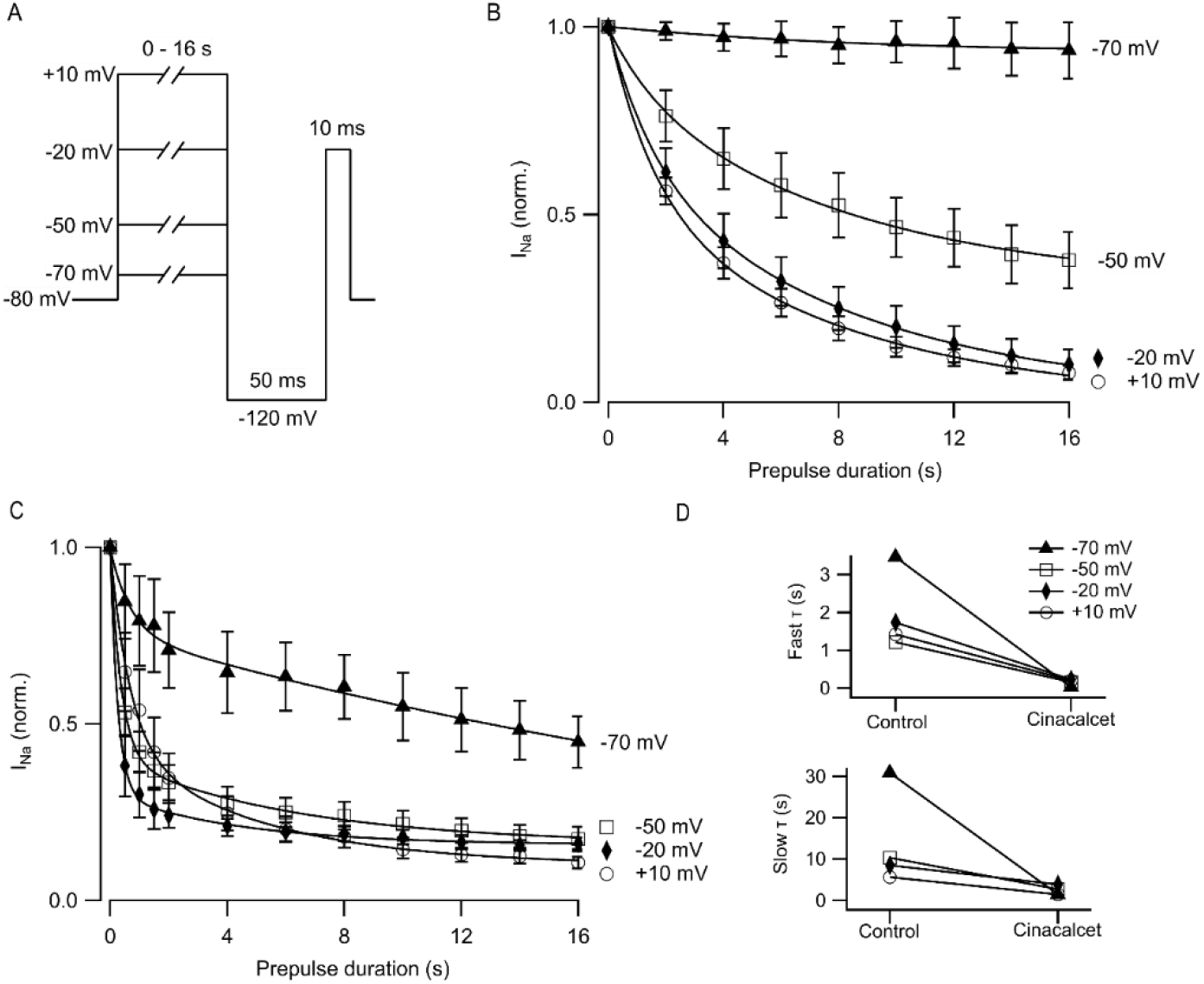
Cinacalcet affinity does not correlate with development of slow inactivation. (A) Voltage protocol used to show development of slow inactivation with increased time at various voltages. Measurements were taken by applying a variable time step to either −70, −50, −20, or +10 mV from a holding potential of −80 mV, and delivering a test pulse to −20 mV following a 50 ms recovery period at −120 mV. (B) Time course of development of slow inactivation in control conditions, with conditioning pulses as in A from −70 (filled triangle; n = 5), −50 (open square; n = 5), −20 (filled diamond; n = 5), or +10 mV (open circle; n = 5) fit to a double exponential. Currents were normalized to the first current at each voltage when there was no inactivating pulse. (C) Time course of development of slow inactivation following complete inhibition by 5 μM cinacalcet, with conditioning pulses as in A from −70 (filled triangle; n = 6), −50 (open square; n = 6), −20 (filled diamond; n = 6), or +10 mV (open circle; n = 5) fit to a double exponential. Currents were normalized to the first current at each voltage when there was no inactivating pulse. (D) Plot of average fast (top) and slow (bottom) t values in control and following inhibition by cinacalcet.

Figure 4C shows the development of VGSC current block during application of 5 μM cinacalcet at different depolarized voltages. The block by cinacalcet can be compared directly with the rate and voltage dependence of slow inactivation shown in Figure 4B, because the pulse protocols are identical and the individual experiments were paired.Importantly, however,the 50 ms pulse to −120 mV that eliminates fast inactivation in the control setting may not have the same effect during block by cinacalcet. If fast-inactivated VGSCs become bound during the variable length prepulse, then the 50 ms pulse to −120 mV will hypothetically only recover those fast-inactivated channels that are *unbound*. Thus, in control, the test pulse assays channels that are in the slow-inactivated state whereas in the presence of cinacalcet, the test pulse will also assay channels that are bound by X.

At each voltage, the kinetics of block by cinacalcet were substantially faster than the development of slow inactivation. At −70 mV, cinacalcet block developed with a fast time constant of 0.03 s and a slow time constant of 1.46 s whereas slow inactivation developed with a fast time constant of 3.46 s and a slow time constant of 30.86 s (Figure 4D). At −50, −20, and +10 mV, cinacalcet block developed with fast time constants of 0.16, 0.23, and 0.20 s respectively, and slow time constants of 2.7, 3.9, and 1.5 s respectively. Conversely, slow inactivation developed with fast time constants of 1.2, 1.7, and 1.4 s at −50, −20, and +10 mV, respectively, and slow time constants of 10.3, 8.5, and 5.7 s. The results with a prepulse to −70 mV demonstrate most clearly the lack of selective binding of X to the slow-inactivated state. There was minimal slow inactivation even with a prepulse to −70 mV for 16 s, yet there was substantial block by cinacalcet (~55%). These results indicate that cinacalcet does not promote the binding of X exclusively to the slow-inactivated state, as development of block could never be faster than the development of slow inactivation if this were the case.

### Inhibition of VGSC currents by cinacalcet is accelerated at voltages favoring the fast-inactivated state

Previous work has shown that sustained hyperpolarization slowly reverses cinacalcet-mediated block of VGSC currents (Mattheisen *et al*., 2018). We tested if holding potential (−60, −80, or −100 mV) strongly affected the dynamics of recovery from inhibition. A double-pulse protocol (S_1_ and S_2_, each to −20 mV for 10 ms) was used to elicit VGSC currents (I_1_ and I_2_) in control or after complete block by 5 μM cinacalcet (Figure 5A). The ratio I_2_ / I_1Con_ reflected the amplitude of the VGSC current elicited by S_2_ compared to that elicited by the step to −20 mV from V_h_ before drug application. At −80 mV, I_2_ / I_1Con_ recovered fully within 10 ms before cinacalcet application (Figure 5B). Cinacalcet attenuated and slowed the recovery of I_2_ / I_1Con_ substantially, so that it eventually reached between 31 and 53%of the control I_2_ / I_1Con_, depending on V_h_, elicited after 8 s at −120 mV (Figure 5B). The majority of this recovery was well described by a single exponential where *τ* was between 1 and 1.9 s (red curves, Fig. 5B). V_h_ had a much greater effect on the dynamics of recovery of the control VGSC currents than the recovery following block by cinacalcet.

**Fig. 5.**
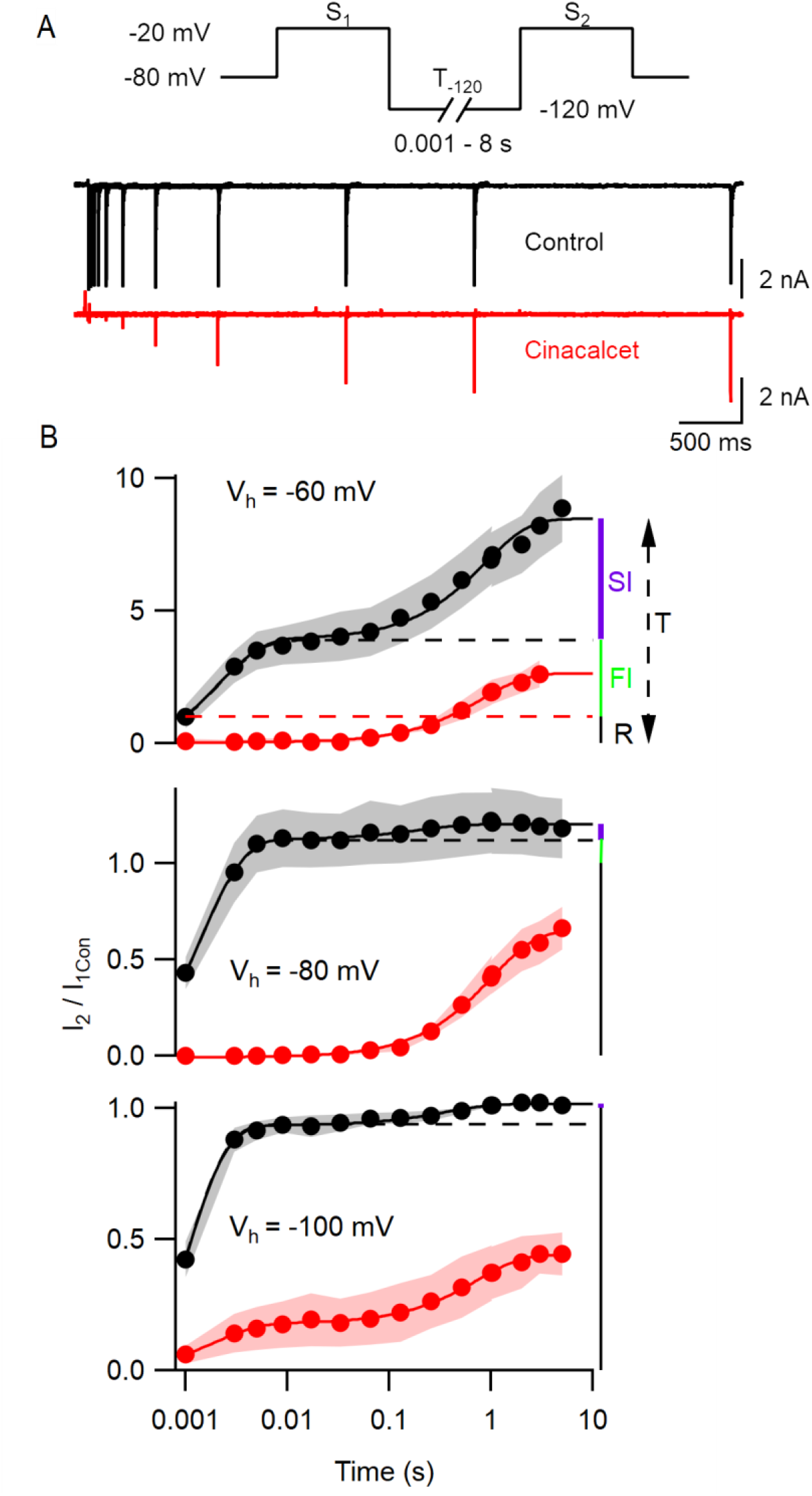
Cinacalcet block is reversible by prolonged hyperpolarization. (A) Superimposed representative traces from a double pulse protocol (S1 and S2) used to elicit VGSC currents in control (top, black) or after complete block by 5 μM cinacalcet (bottom, red) in the whole cell from a holding potential of −80 mV. Test pulses S1 and S2 are 10 ms in length and separated by a variable-length recovery period at −120 mV. (B) Graph showing double-exponential increase in VGSC current amplitude with increased time at −120 mV in control conditions (black) (T1 = 2.08 ± 0.49 ms, T2 = 868.8 ± 119 ms; n = 4) and single-exponential recovery of VGSC current after full inhibition with 5 μM cinacalcet (red) with increased time at –120 mV (T = 816.7 ± 51 ms; n = 4); from a holding potential of −60 mV (top panel) in the whole cell. Graph showing double-exponential increase in VGSC current amplitude with increased time at −120 mV in control conditions (black) (T1 = 1.39 ± 0.09 ms, T2 = 206.1 ± 62.05 ms; n = 4) and single-exponential recovery of VGSC current after full inhibition with 5 μM cinacalcet (red) with increased time at −120 mV (T = 1,018 ± 54.3 ms; n = 5); from a holding potential of −80 mV in the whole cell (middle panel). Graph showing double-exponential increase in VGSC current amplitude with increased time at −120 mV in control conditions (black) (T1 = 0.919 ± 0.059 ms, T2 = 365.4 ± 70.5 ms; n = 4) and double-exponential recovery of VGSC current after full inhibition with 5 μM cinacalcet (red) with increased time at −120 mV (T1 = 1.9251 ± 0.316 ms, T2 = 733.39 ± 46.4 ms; n = 4); from a holding potential of −100 mV in the whole cell (bottom panel). Currents are normalized to current elicited by step from each respective holding potential to −20 mV before cinacalcet addition and to equivalent IV step, in cinacalcet and control traces, respectively.

Models that describe complex VGSC function indicate the channels can occupy multiple fast and slow inactivation states (Ulbricht, 2005; Milescu *et al*., 2010; Goldfarb, 2012; Cervenka *et al*., 2018).The indirect mechanism of block by cinacalcet adds further complexity. To reduce the number of parameters we described the action of cinacalcet using a simpler model with only single fast and slow inactivation states. The two inactive states were were idenitified by the two exponential phases of recovery of I_2_ / I_1Con_ representing VGSC repriming (Fig. 5B, black circles) in the absence of cinacalcet. The amplitudes of the exponentials and the total I_2_ / I_1Con_ value (T, upper panel Figure 5B) were used to estimate the fraction of VGSCs in resting (R, black), fast-inactivated (FI, green), and slow-inactivated (SI, purple) states. At each holding potential, the resting VGSC current was equal to the value I_1Con_ and corresponded to an I_2_ / I_1Con_ value of unity (red broken line, Figure 5B). Thus, the fraction of VGSCs in the resting state was 1/T. The FI component was defined as any component of I_2_ / I_1Con_ below the asymptote for the faster exponential fit (black broken) above the resting component (I_2_ / I_1Con_ =1, broken red line). The SI component was the difference between the asymptote to the double exponential fit (black solid) and the higher of the asymptote of the faster exponential fit (black broken) or I_2_ / I_1Con_ = 1. The fraction (*F*) of the VGSCs in each state was obtained by dividing each component by T (*F_R_, F_FI_*, and *F_SI_*) at each value of V_h_. Using the law of mass action and the observation that reversal of VGSC block was relatively slow (Fig. 5B and (Mattheisen *et al*., 2018)), the rate of block of VGSCs (dB/dt) by the unidentified blocking molecule was directly proportional to the sum of the products of *F* and *k* for each channel state:

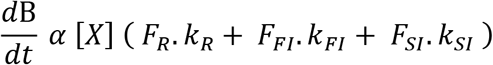

where [X], represent the unknown concentration of blocking molecule, and k the association constants for the three VGSC states. By incorporating a constant, C:

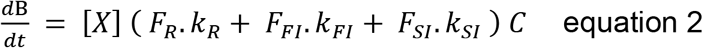

Using equation 2, the voltage-dependent rates of block (1 / *τ*, Figure 1D), and the values of F for each of the states at V_h_ (Table 1) we constructed a simultaneous equation for each of the three holding potentials. These three equations were solved to return the relative association rates for the R, FI, and SI states of 1, 10.1, and 2.3 respectively.

**Table 1:** Parameters used in Model.

| **V<sub>h</sub> (mV)** | **R** | **FI** | **SI** | **1 / $\tau$ (s<sup>-1</sup>)** |
| --- | --- | --- | --- | --- |
| -60 | 0.118 | 0.342 | 0.540 | 0.0096 |
| -80 | 0.832 | 0.100 | 0.069 | 0.0040 |
| -100 | 0.985 | 0 | 0.015 | 0.0020 |

### Cinacalcet state-dependent inhibition of VGSC currents confers non-linear spike block

The increased affinity of cinacalcet activated blocking molecules for the fast-inactivated VGSC state adds a layer of complexity to the effect of this sodium conductance inhibitor on action potential generation. We tested how cinacalcet impacted excitability in neocortical neurons at membrane potentials between −60 and −80 mV. Whole-cell current clamp recordings were performed in the presence of (in μM) 10 CNQX, 10 Gabazine, and 50 APV to prevent confounding by the actions of cinacalcet on synaptic transmission (Vyleta & Smith, 2011). Action potentials were elicited by a series of one second current injections (10-70 pA) in neurons current-clamped at −80 mV (Figure 6A, left panel). The recording mode was switched to voltage clamp and 2 μM cinacalcet applied at −80 mV. After 5 min, the recording mode was switched back to current clamp and depolarizing pulses injected from −80 mV. Cinacalcet was increased to 5 μM and the process repeated as before (Figure 6A, left column). Action potential number decreased at each current injection as cinacalcet concentration was increased and these effects were larger in separate recordings where the membrane potential was held at −70 or −60 mV (Figure 6A, B). A similar concentration-dependence of cinacalcet on excitability was also observed when the sums of action potentials generated at each holding potential were compared (Figure 6C).

**Fig. 6.**
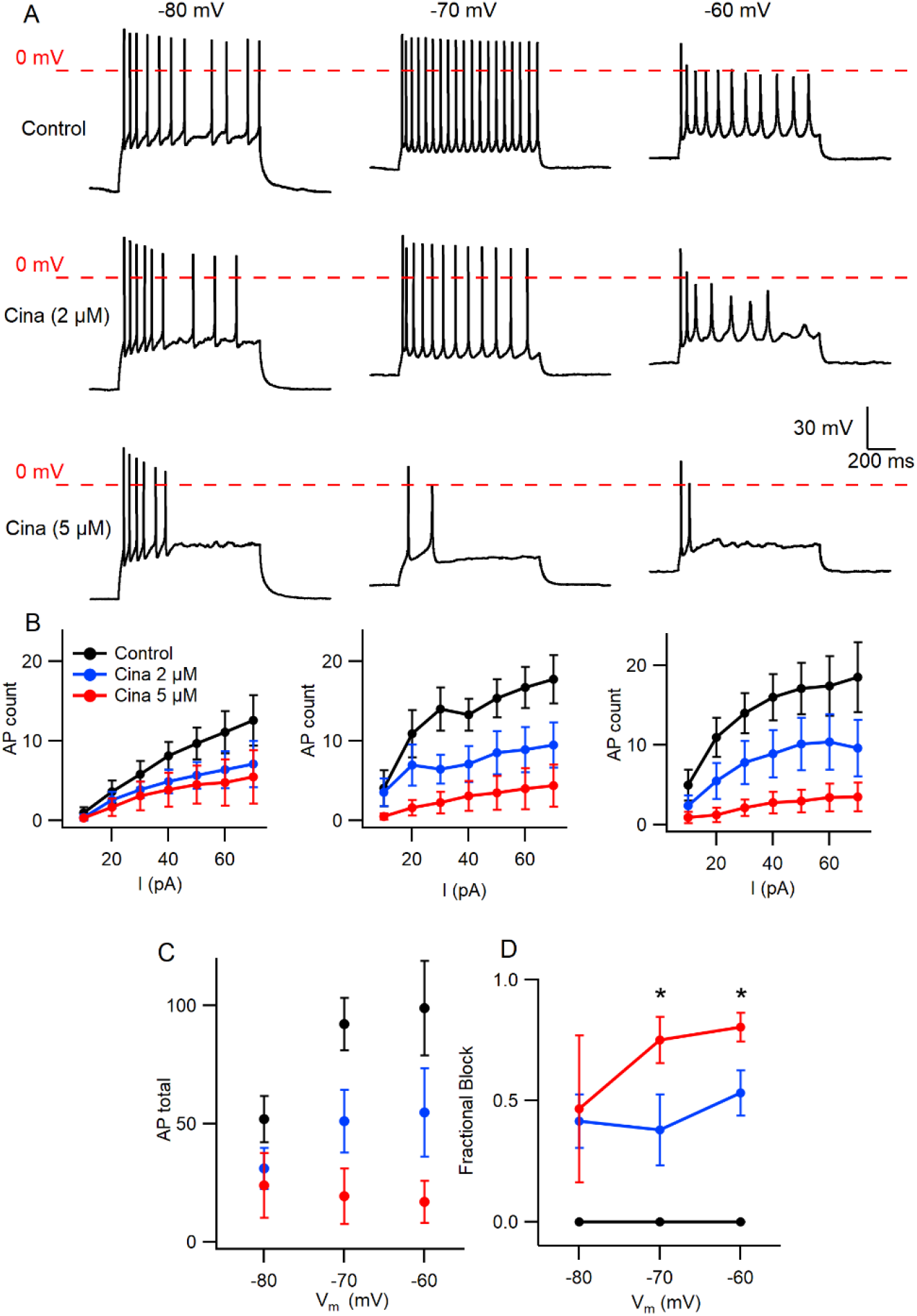
Voltage-dependent inhibition of spike generation by cinacalcet. (A) Exemplar traces showing voltage- and concentration-dependent inhibition by cinacalcet. Three whole-cell current clamp recordings with holding potentials at −80 mV, −70 mV and −60 mV. Measurements were taken at 70 pA before and after cinacalcet was applied for 5 min at incremental doses of 2 μM and 5 μM. (B) Action potential (AP) count following serial current injections from 10 pA to 70 pA for 1 s at holding potentials of −80 mV, −70 mV and −60 mV (from left to right) showing decrease in average AP number with increasing cinacalcet dose from control (black), 2 μM (blue) and 5 μM (red) as well as increased separation of the curves as more depolarized holding potentials. (C) Plot demonstrating a comparison of the cumulative number of APs as seen in B. (D) Fractional Block of AP during maximal current injection of 70 pA at each holding potential. The fractional block with 5 μM cinacalcet increased from 46% to 75% to 80% with depolarization of holding potential from −80 mV to −70 mV to −60 mV. In addition the dose-dependence of the fractional block increased in a non-linear manner with no difference at −80 mV but showed a significant increase in block at −70 mV and −60 mV with increasing cinacalcet dose (mean fractional block 42 ± 11 % vs. 47 ± 30 % at −80 mV, 38 ± 15 % vs. 75 ± 10 % at −70 mV, and 53 ± 9 % vs. 80 ± 6 % at −60 mV at 2 μM and 5 μM; n=10-11 for all groups; * p <0.05 by paired t-test).

Using the number of spikes elicited by the 70 pA current injection we examined how the voltage-dependence of cinacalcet inhibition of sodium conductances (Figure 1) impacted action potential generation (Figure 6D). While the average fractional block of action potentials was the same for 2 and 5 μM cinacalcet at −80 mV (42 ± 11 % vs. 47 ± 30 %), the relative block of the two concentrations increased substantially at −70 mV (38 ± 15 % vs. 75 ± 10 %). Further depolarization to –60 mV reduced the difference in fractional block between the two concentrations of cinacalcet (53 ± 9 % vs. 80 ± 6 %).

Overall, stronger depolarizations increased the potency of action potential block by cinacalcet.

## Discussion

Cinacalcet inhibits VGSC currents strongly in the vast majority of neocortical and hippocampal neurons (Mattheisen *et al*., 2018). Characterizating the mechanism of this prevalent and high-efficacy inhibition will help determine its role in regulating cortical excitability. Here we demonstrate how cinacalcet inhibits the VGSC current by activating a downstream blocking molecule that preferentially binds to the fast-inactivated state, how this stabilizes the inactivated states, and how this impacts neuronal excitability in a non-linear manner.

In our investigation of inactivated VGSC state preference, we used voltage protocols designed to evoke and study the fast- and slow-inactivated states, and the ways in which these states are shifted by the addition of cinacalcet (Figures 2 and 3). Each inactivation curve was shifted in the hyperpolarizing direction by the addition of cinacalcet, indicating stabilization of the inactivated state, and the addition of cinacalcet greatly enhanced the proportion of channels recovering slowly (Figure 3). We found that fast and slow inactivation were both shifted significantly after block with 5 μM cinacalcet by −33 and −38 mV respectively (Figure 2 and 3). The voltage dependence of VGSC current inhibition by cinacalcet is characteristic of many sodium channel inhibitors (SCIs), and can be understood by the modulated receptor model (Hille, 2001) which describes how preferential binding to a specific channel state disturbs the dynamic equilibrium, causing a counteracting shift and new position of equilibrium. The principle of microscopic reversibility ensures that tighter binding to the inactivated state by the blocking molecule results in a greater fraction of the unblocked channels residing in the inactive state at that voltage, = corresponding to a hyperpolarizing shift in V_0.5_ (Hille, 1977, 1978; Bean *et al*., 1983). To further distinguish between the possibilities of selective binding to the slow-inactivated state and slow binding to the fast-inactivated state, we used a protocol to investigate the kinetics of slow inactivation as previously described (Kuo & Bean, 1994) (Figure 4). The results of this experiment argue against the possibility of selective binding to the slow-inactivated state, as the development of block by cinacalcet proceeds at a rate faster than the development of slow inactivation. However, interpretation of data obtained with this approach may not be so straight forward. For example, with this approach it is difficult to distinguish VGSC recovery from block if the channels are in the slow-inactivated or fast-inactivated states when the dissociation of the blocking molecule is relatively slow (Karoly *et al*., 2010). Since cinacalcet acts indirectly to block VGSCs it seems unlikely that external concentrations of cinacalcet are linearly related to the concentration of the downstream blocking molecule. In the absence of the information about the effects of changing the concentration of the blocking molecule we estimated the relative affinity of the blocking molecule for the R, FI, and SI states using equation 2. The normalized inward currents elicited by the double pulse protocol before and after perfusion of cinacalcet were plotted at three separate holding potentials (Fig 5). We used a rate constant derived from the median rates of block at these holding potentials and utilized assumptions such as relatively slow off rate for the blocking molecule, a relatively abrupt increase and stable concentration of blocking molecule following the application of cinacalcet, that all VGSC isotypes respond similarly to cinacalcet application, and that interconversions between channel states are relatively rapid compared to the blocking molecule actions. Based on this equation, the estimated relative affinities for the various states were FI:SI:R in the ratio 10.1 : 2.3 : 1. The accuracy of these predictions will be tested as other components in this pathway are identified and can be incorporated into the model.

As mentioned above, CaSR is not the GPCR transducing the action of cinacalcet and the identity of the cinacalcet target remains unclear. CaSR interacts with the GABA**_B_** receptor in some cells (Chang *et al*., 2020) but this receptor did not contribute to inhibition of VGSC currents by cinacalcet (Mattheisen *et al*., 2018). The muscarinic acetylcholine receptor M1, dopamine receptor D1, and metabotropic receptor mGluR1 have all been identified as GPCRs that can regulate VGSC currents in the cortex via PKA or PKC (Cantrell *et al*., 1996; Cantrell *et al*., 1997; Carlier *et al*., 2006) but agonists and antagonists operating via these receptors did not modulate VGSC currents that were sensitive to cinacalcet (Mattheisen *et al*., 2018). Nor did a wide range of blockers of PKA and PKC, indicating that cinacalcet is operating via a different mechanism than those utilized by acetylcholine, dopamine, and glutamate. The pathway utilized by cinacalcet to modulate VGSC currents also has a higher efficacy and slower timecourse than those activated by acetylcholine, dopamine, and glutamate. For instance the rates of block and unblock by cinacalcet are more than an order of magnitude slower than muscarinic agonists (Figure 1) (Cantrell *et al*., 1996; Mattheisen *et al*., 2018). Consequently, activation of the pathway used by cinacalcet will provide a much slower pattern of modulation of neuronal excitability. The shift in gating of slow and fast inactivation by cinacalcet was more than −30 mV (Figure 2 and 3) whereas gating was unaffected by dopamine agonists (Cantrell *et al*., 1997). This higher voltage-dependence of block by cinacalcet will result in cinacalcet impacting excitable cells that are depolarized much more than those that are hyperpolarized. Comparable differences in the voltage dependence of ion channel blockers has been shown to result in enormous differences in tissue-specific potency. A vivid example is provided by dihydropyridines where at therapeutic levels the L-type cardiac calcium channels are unaffected whereas those in relatively depolarized smooth muscle cells are blocked (Bean, 1984).

We have determined that the action of cinacalcet is highly state dependent and that VGSC block is favored especially when the fast-inactivated state is more preponderant. This state dependence of cinacalcet’s effect, manifested as use-dependent block (Mattheisen *et al*., 2018) and strong dependence of action potential block on the neuronal membrane potential. The prevalence and efficacy of the signaling pathway by which cinacalcet inhibits VGSC positions it to inhibit neocortical neuronal excitability nonuniformly, with major impact on active circuits containing more depolarized neurons. Combined with the unusual pattern of use-dependence (Mattheisen et al, 2018) and the large difference in rate of block over the resting membrane potential range (Figure 1,6) it is likely that cinacalcet will alter cell excitability differently to many other SCIs at the organism level. Further identification of the molecular components of the pathway will facilitate the development of analogous ligands that may avoid co-stimulation of the CaSR and be useful additions to the armamentarium of therapeutic SCIs.

## Additional information section

All data supporting the results presented in the manuscript are included in the manuscript figures.

All authors disclose that they have no competing interests/conflicts of interest.

## Authorship contributions

Participated in research design: Smith, Lindner, Rajayer.

Conducted experiments: Lindner, Rajayer

Performed data analysis: Lindner, Rajayer, Smith

Wrote or contributed to the writing of the manuscript: Lindner, Smith Rajayer

## Funding

This work was supported by grants awarded by NIGMS (GM134110) and U.S.

Department of Veterans Affairs (BX002547) to SMS and MRF (ECI1021099) and NIHLB (T32HL083808) to SRR. The contents do not represent the views of the U.S.

Department of Veterans Affairs or the United States Government.

## Acknowledgements

We thank Mr Luke Steiger, Ms Maya Feldthouse, and Dr Timur Tsintsadze for helpful discussions.

